# Time-resolved multi-omics reveals constitutive defense preparedness and early responses to soybean cyst nematode in wild soybean

**DOI:** 10.64898/2026.11.25.769241

**Authors:** Ping Yates, Zhengzhi Tan, Sibesh Maharjan, Yatendra Singh, Yiqing Wang, M. Shahid. Mukhtar, Sixue Chen, Bao-Hua Song

## Abstract

Soybean (*Glycine max*), a versatile cash crop, is the largest global source of plant-based protein and second-largest source of vegetable oil. However, soybean cyst nematode (SCN; *Heterodera glycines*) is the most economically damaging pest of soybean worldwide, causing substantial yield losses annually. Currently, the effectiveness of widely deployed genetic resistance in soybean cultivars is increasingly challenged by evolving SCN populations; therefore, identifying new sources and mechanisms of SCN resistance is critical. Crop wild relatives, the ancestral or closely related wild species of domesticated crops, are rich sources of genetic diversity encompassing various beneficial traits, including pathogen resistance. In this study, we applied a comparative, multi-omics approach to examine the molecular mechanisms underlying SCN resistance in wild soybean (*Glycine soja*). We integrated transcriptomic, metabolomic, and proteomic data from two wild soybean genotypes representing contrasting responses to SCN (S54, resistant; S67, susceptible) at early- and late-stage infection. Our results reveal distinct constitutive molecular profiles and contrasting genotype-specific responses to SCN infection. The resistant genotype S54 exhibited a stronger early transcriptional response, particularly involving defense signaling and stress-responsive processes, whereas the susceptible genotype S67 showed a delayed response. Gene co-expression and integrative multi-omics analyses identified coordinated molecular networks associated with resistance, including defense-related genes and metabolic features. Together, these findings suggest that SCN resistance in wild soybean involves both pre-existing molecular differences and early activation of defense-associated responses. Our work identifies candidate genes, metabolites, and pathways for future functional testing. Ultimately, these findings provide an integrated multi-omics view of SCN resistance mechanisms in wild soybean, extending our understanding beyond commonly studied cultivar-based resistance and offering potential targets for soybean improvement.

## INTRODUCTION

Soybean (*Glycine max* (L.) Merr.) is a versatile cash crop used for human food, animal feed, biofuel, and other industrial applications while also serving as the largest global source of plant-based protein and second-largest source of vegetable oil. Soybean cyst nematode (SCN; *Heterodera glycines* Ichinohe) is a cyst-forming plant pathogen that parasitizes soybean roots to complete its life cycle. SCN is recognized as the most destructive pathogen affecting soybean production, resulting in substantial yield and economic losses (e.g., >$1.5 billion annually in the United States alone). Modern agricultural practices have facilitated SCN’s geographic expansion into new regions, and once established, it is extremely difficult to eradicate (Arjoune et al., 2022; Bandara et al., 2020). Current SCN management strategies heavily rely on the use of resistant cultivars (e.g., PI88788 and Peking) and traditional agricultural practices (e.g., crop rotation). Specifically, approximately 95% of SCN-resistant soybean cultivars have historically relied on resistance derived from PI 88788, particularly the *Rhg1-b* locus (Bent, 2022; Miraeiz et al., 2026). With this sustained selective pressure, SCN populations have rapidly adapted to overcome the most common resistant cultivars as evidenced by reports of surging SCN reproductive rates (up to 90%) in the last few years (Tylka, 2022). These challenging circumstances underscore the urgent need to diversify our genetic sources of SCN resistance and move beyond our reliance on resistance mechanisms derived from single genetic sources.

Crop wild relatives (CWRs), the ancestral or closely related wild species of domesticated crops, provide rich sources of genetic diversity encompassing a broad range of agronomically-beneficial traits, including pathogen resistance (Bohra et al., 2022; Harlan, 1976). Despite their indispensable historical role, CWRs have been largely neglected and underutilized in modern crop breeding and improvement programs. CWRs, such as wild soybean (*Glycine soja* Sieb. et Zucc.), harbor a wealth of genetically diverse alleles that could compensate for losses in genetic diversity associated with soybean domestication and modern breeding (Kim et al., 2023; Li et al., 2024; Li et al., 2014; Zhang et al., 2016). Although numerous SCN-resistant accessions have been identified, very few genes and resistance loci have been characterized at the molecular level in soybean accessions other than the resistant cultivars Peking and PI88788 (Miraeiz *et al*., 2026). Among identified resistant accessions, wild soybean genotypes have demonstrated their potential as emerging sources of SCN resistance, but compared to cultivated soybean, research on their underlying resistance mechanisms remains limited (Grunwald et al., 2022; Kofsky et al., 2021; Lakhssassi et al., 2025; Miraeiz et al., 2020; Zhang *et al*., 2016). Beyond providing alternative resistance alleles for crop improvement, wild soybean offers opportunities to investigate naturally evolved defense mechanisms that may differ from those extensively characterized in cultivated soybean.

Over the past decade, emerging approaches, including genome editing, systems biology, and multi-omics integration, have unlocked new opportunities to study host-pathogen interactions and identify new sources of resistance (Samal et al., 2026). Typically, pathogen-induced responses have been characterized at the single-omics level, most commonly using transcriptomics or metabolomics (Huang et al., 2022; Tian et al., 2025). Although these insights are valuable, this approach provides a limited understanding, as patterns across individual omics layers do not always predictably align. Integrating transcriptomic, proteomic, and metabolomic data provides an opportunity to investigate how constitutive molecular differences and pathogen-induced responses are coordinated across regulatory and metabolic networks, potentially revealing resistance-associated features that may be overlooked by individual omics analyses (Kumar et al., 2025). Together, multi-omics data paired with rigorous computational tools may uncover interconnected molecular mechanisms underlying SCN resistance and accelerate the discovery of molecular targets for improving SCN-resistant cultivars.

Although soybean-SCN interactions have been extensively studied, our mechanistic understanding of SCN resistance comes primarily from resistant cultivars, mainly PI88788 and Peking (Bent, 2022). Thus far, the molecular strategies underlying naturally evolved SCN resistance in wild soybean remain poorly understood. Furthermore, most investigations have studied individual molecular layers, providing only fragmented views of resistant responses, meanwhile, reports integrating two or more omics layers to compare susceptible and resistant soybean responses to SCN have been rare (Shi et al., 2021). In particular, it remains unclear how constitutive molecular differences and infection-induced responses contribute to resistance. How these responses change over the course of infection, and how resistance-associated features are connected across transcriptional, proteomic, and metabolic networks. Addressing these knowledge gaps may reveal molecular strategies associated with SCN resistance in wild soybean that have received limited attention in cultivated soybean systems.

To address these research gaps, we applied a comparative, multi-omics approach integrating transcriptomic, proteomic, and metabolomic data from two wild soybean genotypes with contrasting responses to SCN (S54, resistant; S67, susceptible) at both early-and late-stage infection. Here, we investigate three fundamental questions: (1) What constitutive and SCN-induced molecular features distinguish resistant S54 from susceptible S67? (2) How do the timing and coordination of transcriptional, proteomic, and metabolomic responses to SCN infection differ between resistant and susceptible genotypes? (3) Which features are shared across co-expression and multi-omics networks and may contribute to SCN resistance in wild soybean? By integrating molecular responses across multiple omics layers and infection stages, we aimed to identify resistance-associated features and candidate mechanisms that may expand our understanding of naturally evolved SCN resistance beyond the extensively studied cultivated soybean systems.

## METHODS

### Study Materials and Experimental Setup

Wild soybean accessions, S54 (PI 424093) and S67 (PI 468396 B), were obtained through GRIN-Global (USDA National Plant Germplasm System). S54 and S67 were previously identified as resistant (R) and moderately susceptible (MS) to SCN Hg Type 1.2.5.7 (Race 2). Race 2 SCN was sourced directly from Song Laboratory stocks maintained on Williams 82 under greenhouse conditions. Our experimental design included 2 genotypes (S54, S67), 2 treatments (SCN-infected, control), and 2 timepoints (3 and 8 days post-inoculation, dpi) (**Fig. 1**). Seeds were disinfected with 5% sodium hypochlorite for 3 minutes, rinsed with deionized water, and scarified using a scalpel prior to germination. Plants were grown in sterile, sand-filled cone-tainers (Greenhouse Megastore, IL, USA) under controlled chamber conditions (27°C, 16h light cycle) with daily watering. Plant positions were rotated every two days to minimize spatial effects.

**Figure 1.**
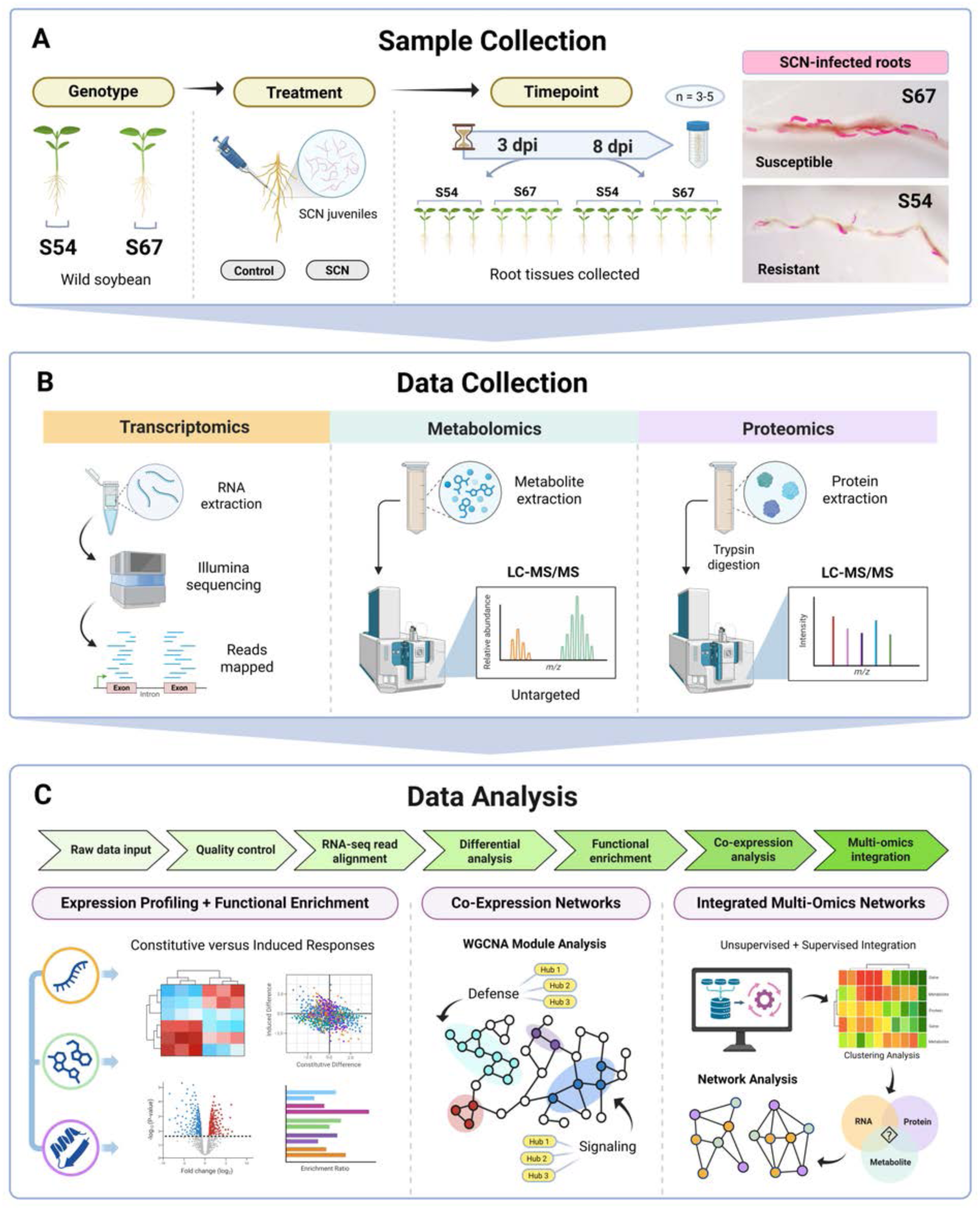
**Schematic of project workflow**. (A) Sample collection of resistant (S54) and susceptible (S67) wild soybean genotypes infected with SCN at 3 and 8 dpi timepoints. Representative microscope images of acid fuchsin-stained SCN juveniles in 8 dpi roots are shown for both genotypes. (B) Data collection from root tissue samples. Transcriptomics (RNA-seq; n = 3 per condition). Metabolomics and proteomics (LC-MS/MS; n = 3-5 per condition). (C) Overview of data analysis approach for single-omics and multi-omics. Graphic generated in BioRender.

### Nematode Collection and Inoculation

Race 2 SCN eggs were collected from Song laboratory stocks using standard nested, sieve-based extraction methods, 70% sucrose flotation, and centrifugation (Zhang et al., 2017). Eggs were disinfected with 0.5% sodium hypochlorite for ∼3 minutes and thoroughly rinsed with water. Eggs were hatched *in vitro* by incubating eggs for 3 days at 27°C in darkness using the Baermann tray technique supplemented by 10-20 mM ZnSO_4_. After 3 days, J2-stage juveniles were collected in a 25 µm sieve, examined for viability, and counted under a stereomicroscope. Eight-day old seedlings were inoculated with ∼2,000 juveniles (J2) per plant or mock-inoculated with water (control groups). Inoculum was added through two holes adjacent to the roots, with juveniles maintained in suspension by continuous stirring with a magnetic rod to ensure uniform distribution.

### Tissue Collection

Fresh root tissue was collected from control and SCN-infected plants at 3 and 8 days post inoculation (dpi). Each sample consisted of pooled tissue from 3–6 plants. Samples were immediately flash-frozen in liquid nitrogen and stored at -80°C. Frozen root tissues were homogenized using sterile, steel beads with a Bullet Blender 5E Gold+ tissue homogenizer (Next Advance Inc., Troy, New York, USA) for subsequent RNA, metabolite, and protein extraction.

### RNA Sequencing

RNAseq libraries were prepared from extracted RNA samples (3 replicates per experimental group) and sequenced on an Illumina NextSeq 2000 short-read sequencer (paired-end, P4 300-cycle kit) at the University of Delaware DNA Sequencing and Genotyping Center (Newark, Delaware, USA). Illumina library quality was assessed using Qubit, Fragment Analysis, and ddPCR.

### Metabolite and Protein Extraction and Processing

Root samples were processed in Sixue Chen’s lab at the University of Mississippi. Tissues were freeze-dried in a FreeZone Triad Freeze Dryer (Labconco Corporation, Kansas City, USA) for 48 hours at -50°C under a vacuum of 0.012 mbar. Dry weight samples (15 mg) were stored in a desiccator at 4°C for further analysis. Sample tubes containing one 6-mm stainless steel grinding ball (Cole-Parmer, Vernon Hills, IL, USA) were immersed in liquid nitrogen and homogenized twice using a Cole-Parmer Spex HG-400 MiniG homogenizer (1000 rpm for 2 min). Metabolite extraction was performed as previously described with minor modifications (Li et al., 2023). Lidocaine and 10-camphorsulphonic acid (10 µL each, 100 µM) were added as internal standards in each sample. Sequential extraction was performed using cold (–20°C) extraction solvents: acetonitrile:isopropanol:water (3:3:2, v/v/v; 800 µL), acetonitrile:water (1:1, v/v; 600 µL), and 80% methanol (600 µL). Samples were vortexed (10 min), sonicated in ice-water (2 min), and centrifuged at 15,000 × g for 10 min at 4°C. Supernatants were collected and lyophilized to dryness. The remaining pellets were used for protein extraction as previously described (Guan et al., 2025). The pellets were solubilized in 95 μL of protein solubilization solution by vortexing (30 min at 20°C), sonicated in ice-water (2 min, repeated 3 times), and centrifuged (15,000 × g for 20 min at 4°C) to remove debris. Supernatants were collected and stored at −20°C. For protein cleanup, 200 μL of 100% methanol was added to supernatants (50 μg of protein) and vortexed for 30 s. Chloroform (50 μL) and ddH2O (150 μL) were added and solutions were vortexed for 30 s and centrifuged at 12,000 × g at 4°C for 10 min. After discarding the upper chloroform phase, 150 μL of 100% methanol was added to the lower phase and centrifuged at 15,000 × g and 4°C for 10 min. The pellet was lyophilized to dryness.

### Protein Quantification and In-Solution Trypsin Digestion

Protein concentration was determined using a modified CB-X Protein Assay (G-Biosciences, Geno Technology Inc., St. Louis, MO, USA). Distilled water (8 μL) was added to protein extracts (2 μL), followed by addition of pre-chilled CB-X reagent (1 mL). The mixture was vortexed (10 min), sonicated in ice-water (6 min), and centrifuged (15,000 × g for 6 min at 4°C) before removing the supernatant. Pellets were solubilized by adding 50 μL of CB-X Solubilization Buffer-I and Buffer-II and vortexed until complete dissolution. The remaining steps followed the manufacturer’s instructions with absorbance measurements taken at 595 nm on a Synergy LX Multi-Mode Reader (BioTek Instruments, Inc., Winooski, VT, USA).

Protein samples were reduced and alkylated by adding dithiothreitol (3 μL, 10 mM) and chloroacetamide (11 μL, 500 mM), and incubated (room temperature, 1 h). Reaction volumes were adjusted to 500 μL with 100 mM NH₄HCO₃ (Tan et al., 2025). Trypsin digestion (1:100 ratio, trypsin:protein, w/w) was carried out overnight (∼16 h, 37°C). Digested peptides were lyophilized to dryness using a CentriVap 7812010 Mobile System (Labconco, USA). Resulting pellets were resuspended in 15 μL of 0.1% formic acid (FA), vortexed (10 min), sonicated (ice-water, 5 min), centrifuged (15,000 × g for 10 min at 4°C), and transferred to vials for mass spectrometry (MS) analysis.

### Liquid Chromatography Tandem MS (LC-MS/MS) Based Proteomics

Peptide samples (3.5 μL) were analyzed using a ZenoTOF 7600 system (SCIEX, USA) coupled to a nanoACQUITY LC system (Waters, USA). Chromatographic separation was performed with a trap column (nanoEase M/Z Symmetry C18; 5 µm, 180 µm i.d. × 20 mm, 100 Å) and an analytical column (nanoEase M/Z HSS C18 T3; 1.8 µm, 75 µm i.d. × 150 mm, 100 Å) in a 40°C column oven, with a constant flow rate of 500 nL min⁻¹ over a 95 min run time.

Mobile phases consisted of 0.1% FA in water (A) and 0.1% FA in acetonitrile (B). Gradient elution increased linearly from 1% B to 30% B over 60 min, stepped to 40% B at 70 min, ramped to 80% B at 72 min, maintained until 78 min for column washing, before re-equilibration to 1% B at 80 min, and held until 95 min. MS data were acquired in positive-ion mode using data-independent acquisition (DIA) in SWATH mode and Zeno pulsing. Ion source parameters were set to 4500 V spray voltage, 150°C source temperature, 25 curtain gas, ion source gas 1 (12 psi) and 2 (45 psi). TOF-MS scans covered an m/z range of 400–1500 with an accumulation time of 50 ms, followed by MS/MS acquisition across an m/z range of 140–1500 with an accumulation time of 20 ms using collision-induced dissociation (CID). SWATH acquisition was used with variable precursor isolation windows (m/z ∼400–900) and optimized rolling collision energy. Total cycle time was 2.18 s. Data acquisition was controlled using SCIEX OS software (v3.4.4).

Raw data was processed using DIA-NN (v1.8.1) with default high-precision settings for neural network–based peptide identification and quantification, interference correction, and cross-run normalization. Low-confidence identifications and interfering precursors were automatically removed. Peptide- and protein-level false discovery rates were controlled at 1% using a target-decoy method. DIA-NN aligned retention times and normalized intensities across runs, and protein levels were estimated from total peptide intensities after interference and normalization correction.

### Untargeted LC-MS/MS Metabolomics

Metabolite samples were solubilized in 0.1% FA by vortexing (2 min), sonicated in ice-water (2 min), and centrifuged at 15,000 × g for 15 min at 4°C. Supernatants were analyzed with the LC-ZenoTOF 7600 MS using the same solvent system as for proteomics with a Hypersil GOLD C18 column (2.1×100 mm, 1.9 μm, 175 Å) and a flow rate of 40 µL min⁻¹ in a 40°C chamber. A gradient running over 30 min, starting at 99% A, increased to 95% at 1 min, increased to 5% A at 14 min, held until 18 min, and returned to 99% A by 20 min was used with partial loop injection mode (3 µL per sample).

TOF-MS spectra data was acquired in positive and negative ionization modes over m/z 50–1200, with a duration of 0.1 s. The OptiFlow microflow source was at 350°C with 35 psi curtain gas, gas 1 (30 psi), and gas 2 (35 psi). Spray voltages (+4500 V/−4300 V) and declustering potentials (+40 V/-40 V) were used in positive and negative-mode, respectively. The top 30 precursor ions exceeding 100 counts s⁻¹ (cps) were selected after dynamic background subtraction for MS/MS fragmentation using CID at 30 ± 5 V (positive-mode) or −30 ± 5 V (negative-mode), with an accumulation time of 0.01 s, and Zeno trap pulsing threshold of 20,000 cps.

Raw data were processed using MS-DIAL and SCIEX OS Analytics for peak detection, deconvolution, alignment, and normalization. Metabolites were annotated by MS1 and MS/MS spectral matching using MS-DIAL against PlaSMA, FiehnHILIC, GNPS, Mass Bank, MassBankEU, Vaniya-Fiehn_Natural_Product, and VS19 databases and SCIEX OS Analytics against a published CIeaD phytochemical spectral library (Singh et al., 2026). Spectra were collected in centroid mode with signal intensity (>500 units) and signal-to-noise ratio (S/N > 5) criteria (Singh *et al*., 2026). Positive- and negative-ionization datasets were examined separately and then combined.

### Data Analysis Strategy

The analysis comprised nine modules: (1) Data processing, read alignment, and input preparation: RNA-seq reads were processed and aligned to the reference genome to generate gene-level count matrices. Proteomic and metabolomic quantitative tables were imported, and sample metadata were standardized. (2) Omics-specific quality control and normalization: Missing values, signal distributions, and sample consistency were assessed, followed by filtering, transformation, and normalization appropriate to each omics dataset. (3) Differential analysis: Treatment effects, baseline genotype differences, and genotype-specific treatment responses were evaluated within each timepoint. (4) Functional annotation and enrichment analysis: Genes, proteins, and metabolites were functionally annotated, and differential features were tested for enrichment of biological processes and pathways. (5) Co-expression network analysis: WGCNA was used to identify co-expression modules, assess their associations with experimental traits, and select candidate hub features. (6) Multi-omics sample matching and matrix preparation: Shared samples were matched across datasets, and sample order and metadata were aligned to construct quantitative matrices for joint analysis. (7) Multi-omics integration: MOFA2 was used to characterize variation in multi-omics data, and DIABLO was used to select correlated features across omics datasets associated with sample groups. (8) Post-integration functional enrichment: Features selected by the integration models were tested for enrichment of biological processes and pathways. (9) Candidate gene prioritization: DIABLO feature selection, WGCNA module membership, and expression patterns across experimental groups were combined to prioritize candidate genes.

### Data Processing, Read Alignment, and Input Preparation

Raw and cleaned paired-end RNA-seq reads were assessed using FastQC (v0.12.1), and quality reports were summarized using MultiQC (v1.29). Clean reads were aligned to the *Glycine soja* W05 reference genome (ASM419377v2; GCF_004193775.1) using STAR (v2.7.10b). Alignments were converted to BAM format and sorted by genomic coordinate using SAMtools (v1.10). Gene-level read counts were generated using featureCounts (v2.0.6), the corresponding GTF annotation from NCBI *Glycine soja* Annotation Release 100, and the options -s 0 -p -B -C. Input preparation was performed in R (v4.4.3). Protein-group abundance and metabolite intensity tables were imported using readxl (v1.4.5). RNA-seq sample identifiers were matched to the sample sheet, whereas proteomic and metabolomic metadata were parsed from sample column names. Genotype, treatment, sampling time, and replicate labels were standardized, and feature-by-sample matrices were assembled.

### Omics-Specific Quality Control and Normalization

RNA-seq quality control included library size, detected gene counts, principal component analysis (PCA), and sample-distance analysis. Genes with fewer than 10 total reads across used samples were removed. Variance-stabilizing transformation was performed using DESeq2 (v1.46.0), with genotype, sampling time, and treatment included in the design and blind = FALSE. Protein abundances were transformed as log₂(x + 1). Metabolite zeros were treated as missing values, and sample intensities were normalized by their medians before log₂(x + 1) transformation. For proteomic and metabolomic quality assessment, missing transformed values were replaced with the minimum observed value in the corresponding sample minus one. Missingness, total signal, intensity distributions, and Pearson correlations were examined. Global PCA used the 1,000 most variable features in each dataset.

### Differential Expression Analysis

Differential analyses were performed separately at each timepoint. RNA-seq counts were analyzed using DESeq2 (v1.46.0) and genes with at least 10 total reads across samples were retained. Treatment effects within each genotype were evaluated using treatment-only models. Genotype differences and genotype-by-treatment interactions were assessed using models containing genotype, treatment, and their interaction. Protein-group and metabolite abundances were analyzed using Limma (v3.62.1). Features observed in at least three samples in every group were retained in the model. Metabolite intensities were normalized by sample medians before log₂(x + 1) transformation; protein intensities were log₂(x + 1)-transformed without additional normalization. After filtering, remaining missing values were replaced with the minimum observed transformed value in the corresponding sample minus one. Linear-model contrasts were evaluated using two-sided empirical Bayes moderated t-tests. Limma log₂ fold changes were the fitted contrasts on the transformed and imputed abundance scale. Comparisons included treatment effects within each genotype, baseline genotype differences, and genotype-by-treatment interactions. For **Figure 2**, DEGs required DESeq2-adjusted p < 0.05 and |log₂ fold change| ≥ 1. DEPs and DEMs required unadjusted p < 0.05 (limma p-value) and |log₂ fold change| ≥ 1 in the within-genotype treatment comparisons. Benjamini–Hochberg-adjusted p-values were used for other FDR-based analyses. For baseline genotype and genotype-by-treatment interaction contrasts shown in **Figure 3**, significant features required adjusted p < 0.05 and |log₂ fold change| ≥ 1 for transcripts or ≥ 0.58 for proteins and metabolites.

**Figure 2.**
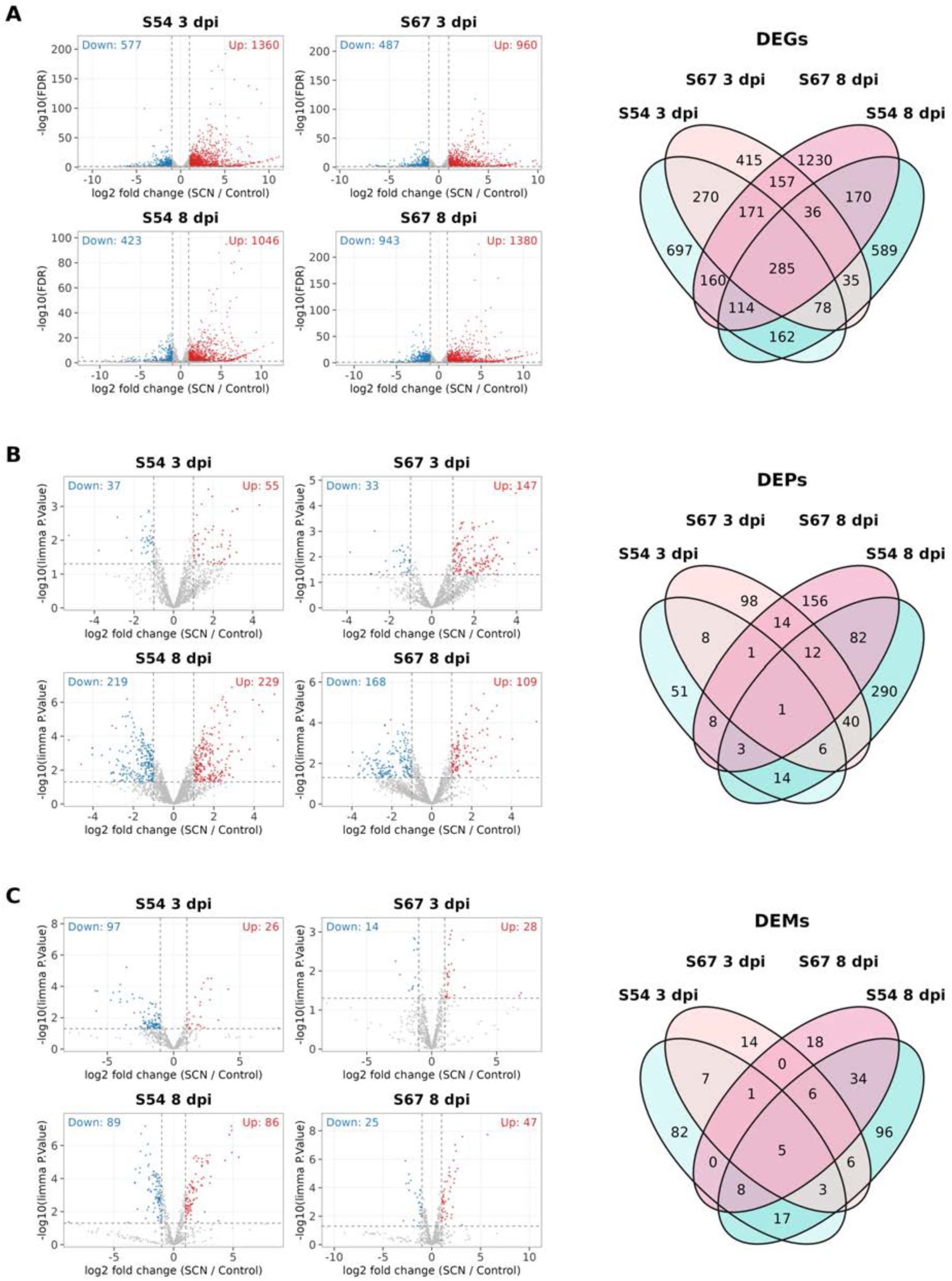
**Differentially expressed features following SCN infection at early (3 dpi) and late-stage (8 dpi) in S54 and S67 samples**. Volcano plots and Venn diagrams show differentially expressed features: (A) genes (DEGs), (B) proteins (DEPs), and (C) metabolites (DEMs). DEGs were defined by DESeq2-adjusted p < 0.05 and |log₂ fold change| ≥ 1. DEPs and DEMs were defined by unadjusted p < 0.05 from limma empirical Bayes moderated t-tests (p-value), together with |log₂ fold change| ≥ 1. All comparisons were made between SCN-treated and control samples within each genotype and timepoint, for RNA-seq (n = 3) and proteomics and metabolomics (n = 4-5). Volcano plots: red, significantly upregulated; blue, significantly downregulated; grey, non-significant features. Venn diagrams summarize feature-set overlaps among the four comparisons, irrespective of direction; each number represents an exclusive intersection, and areas are not proportional to counts.

**Figure 3.**
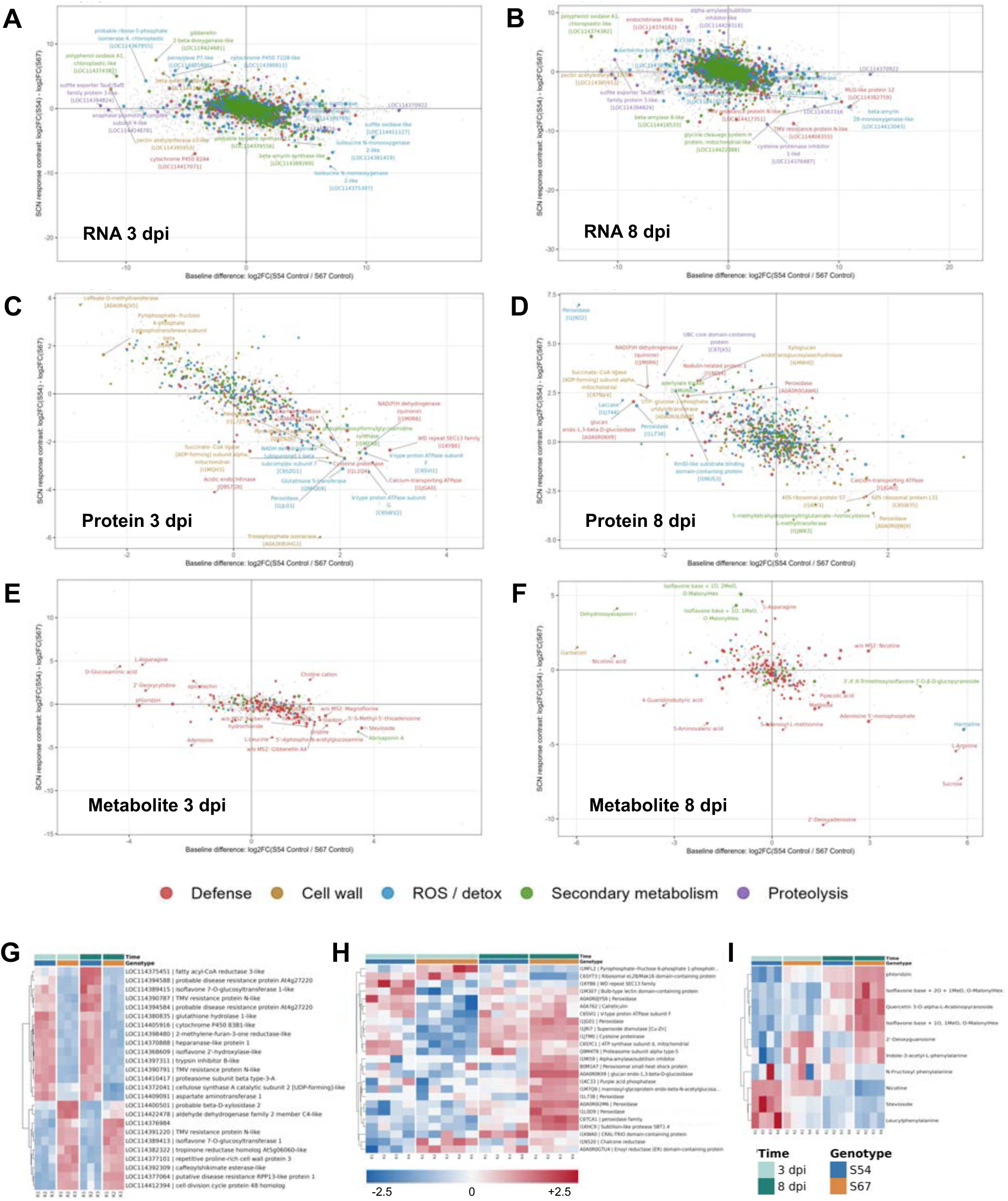
Constitutively expressed features at 3 and 8 dpi in S54 and S67 samples. Quadrant plots show genotype-specific constitutive and differentially (SCN versus control) expressed omics features. RNA features at (A) 3dpi and (B) 8 dpi. Protein features at (C) 3 dpi and (D) 8 dpi. Metabolite features at (E) 3 dpi and (F) 8 dpi. Features are colored based on assigned functional category. Heatmaps show constitutively expressed (z-score) (G) genes, (H) proteins, and (I) metabolites in control samples.

### Functional Annotation and Enrichment Analysis

Gene Ontology annotations were obtained from the NCBI annotation file for the *G. soja* ASM419377v2 assembly. Protein sequences were annotated using eggNOG-mapper (v2.1.13), with the taxonomic scope restricted to Viridiplantae. Annotations from individual protein accessions were combined at the protein-group level. Metabolite features were mapped to KEGG compounds using the local metabolite reference table, and compound-to-pathway relationships were retrieved using KEGGREST (v1.46.0). Metabolite-associated GO terms were inferred indirectly through KEGG pathways and their associated *G. soja* genes. GO term names were obtained using GO.db (v3.20.0) and AnnotationDbi (v1.68.0). Upregulated and downregulated feature sets were analyzed separately using clusterProfiler (v4.14.0), with annotated features tested in the corresponding differential comparison used as the background. For metabolite enrichment, input features were selected using an adjusted p-value < 0.05 and |log₂ fold change| ≥ 0.58. Enrichment p-values were corrected using the Benjamini–Hochberg method, and terms with adjusted p-values ≤ 0.05 were retained.

### Co-Expression Network Analysis

Co-expression networks were constructed using WGCNA (v1.74). For RNA-seq, the 5,000 genes with the highest variance in the variance-stabilized matrix were selected across the 24 samples at each timepoint. A signed network was constructed using biweight midcorrelation and a soft-thresholding power of 13. Modules were identified using blockwiseModules, with a minimum module size of 30 and a merge cut height of 0.25. Proteomic and metabolomic networks used complete-case features, signed Pearson correlations, and soft-thresholding powers of 11 and 12, respectively. Minimum module sizes for proteins and metabolites were 30 and 20, respectively, with a merge cut height of 0.25. Module eigengenes were correlated with experimental traits. Features were ranked within each non-grey module by module membership, defined as the correlation with the corresponding module eigengene, and the top 30 were retained as candidate hub features.

### Multi-Omics Sample Matching and Matrix Preparation

Variance-stabilized RNA-seq counts, log-transformed protein abundances, and normalized log-transformed metabolite intensities were matched using sample identifiers together with sampling time, treatment, and genotype. Samples represented in all three datasets were retained, yielding 22 matched samples, with 11 at each timepoint. Sample order was aligned across matrices. Each feature was centered and scaled using the mean and standard deviation of its observed values, while missing entries were retained. Standardization was performed across all matched samples before the matrices were divided into 3- and 8-dpi datasets.

### Multi-Omics Integration

Unsupervised integration was performed separately at 3 and 8 dpi using MOFA2 [1.16.0]. Up to 2,000 transcriptomic, 2,000 proteomic, and 1,000 metabolomic features were selected by variance. Gaussian models with two latent factors were trained for a maximum of 5,000 iterations, and factor scores, variance explained, and feature weights were extracted. Supervised integration used DIABLO (Anagho-Mattanovich et al., 2026) implemented in mixOmics [6.30.0]. Separate models used treatment or genotype as the response variable. Missing values were replaced with feature-wise medians, and features with insufficient variation were removed. Two-component models were fitted with an inter-block design weight of 0.1. The number of selected features per block and component was tuned over 5, 10, and 20 using three-fold cross-validation repeated five times. Internal predictive performance was evaluated using the same cross-validation settings and maximum-distance classification.

## RESULTS AND DISCUSSION

### Shared SCN responses, but divergent molecular composition and timing

In this study, we applied a multi-omics approach to characterize the response to SCN infection in resistant (S54) and susceptible (S67) wild soybean genotypes at 3 and 8 days post-inoculation (dpi). Among the total identified genes, metabolites, and proteins for each genotype-timepoint subset, we evaluated differentially expressed (SCN versus control) features (DEGs, DEMs, DEPs) (**Fig. 2**).

Across 3 and 8 dpi samples, 2,767 and 3,121 genes in S54 and S67, respectively, met our significance criteria for differential expression (SCN vs. control). At 3 dpi, 1,937 and 1,447 DEGs were detected in S54 and S67, respectively; at 8 dpi, 1,469 and 2,323 DEGs were detected in S54 and S67, respectively. In both genotypes, the majority of DEGs were upregulated at each timepoint, with S54 exhibiting more upregulated DEGs at 3 dpi whereas S67 exhibiting more at 8 dpi (**Fig. 2A**). At the protein level, 516 and 429 proteins in S54 and S67, respectively, met our criteria at one or both time points. At 3 dpi, 92 and 180 DEPs were identified in S54 and S67, respectively; at 8 dpi, 448 and 277 DEPs were identified in S54 and S67, respectively (**Fig. 2B**). For metabolites, 265 and 102 metabolites in S54 and S67, respectively, met the criteria at one or both time points. More DEMs were identified in S54 than in S67 at both timepoints (**Fig. 2C**).

To identify key biological processes underlying these responses, we performed GO and KEGG enrichment analyses of upregulated and downregulated DEGs across each genotype-timepoint subset. Both genotypes showed enrichment of core defense-associated processes, including plant-pathogen interaction, defense and oxidative-stress responses, peroxidase activity, MAPK signaling, and isoflavonoid/phenylpropanoid biosynthesis (**Fig. S1**), indicating that several common pathways are activated following SCN infection. Simultaneously, genotype-specific patterns were also observed: S54 showed prominent enrichment of hormone-mediated signaling, cell wall-related processes, chitinase activity, and specialized metabolic pathways, whereas S67 showed enrichment of transport, cell wall activity, and primary and secondary metabolism processes. Overall, both genotypes exhibited transcriptional reprogramming involving several recognized defense processes, including plant–pathogen interaction, MAPK signaling, oxidative-stress responses, phenylpropanoid metabolism, and isoflavonoid biosynthesis, but differed in the composition and regulation of their molecular responses (**Fig. S1**). The abundance and composition of differential features varied across molecular layers and timepoints. At 3 dpi, more DEGs and DEMs were identified in S54, whereas more proteins were differentially expressed in S67. At 8 dpi, S67 exhibited more differential transcripts, whereas more proteins and metabolites were differentially expressed in S54 (**Fig. 2**). Notably, the contrasting transcriptional patterns between S54 and S67 raise the possibility that effective resistance involves distinct early defense responses, whereas the more extensive transcriptional changes in S67 at 8 dpi may reflect continued infection-associated stress or compensatory responses. These feature counts alone do not establish global differences in genotype-specific treatment effects; however, they reveal interesting genotype-associated and temporal patterns of SCN responses that warrant further investigations. Expanded temporal sampling would help distinguish differences in response timing from differences in response magnitude or persistence.

### Constitutive defense-associated molecular differences distinguish resistant and susceptible wild soybean genotypes

Comparative analysis of transcriptomic, metabolomic, and proteomic profiles revealed distinct constitutive expression patterns between S54 and S67 control samples (**Fig. 3**). Comparisons between control samples identified more genes with significantly higher expression in S54 than in S67, particularly at 3 dpi. A subset of defense-and cell wall-related genes also showed greater constitutive expression in S54 than in S67. These genes were related to pathogen resistance (e.g., TMV resistance protein N-like), metabolic processes (e.g., cytochrome P450 83B1, glutathione hydrolase 1, isoflavone 2′-hydroxylase-like), cell wall modification (e.g., cellulose synthase A), and proteolysis (**Fig. 3G**). Constitutive differences were also observed at the protein level. S54 control samples showed a few more highly expressed proteins than S67 control samples, mainly at 3 dpi. Among selected defense-and cell wall-related proteins, S54 showed higher constitutive abundance of a bulb-type lectin protein, peroxidase, calreticulin, ATP synthase, and proteasome subunit (**Fig. 3H**). Across metabolomic samples, S54 and S67 control samples showed fairly similar overall metabolite profiles. However, a subset of defense- and cell wall-related metabolites showed constitutive differences between S54 and S67, particularly at 3 dpi (**Fig. 3I**). S54 exhibited higher constitutive abundance of metabolites including alkaloids, terpenoids, and leucylphenylalanine than S67. Together, these constitutive molecular differences suggest that resistant (S54) and susceptible (S67) genotypes occupy distinct molecular baseline states under control conditions, which may contribute to the overall effectiveness of their defense response.

Collectively, these constitutively abundant defense-associated features may contribute to structural, biochemical, and molecular preparedness for pathogen challenge, including antimicrobial metabolites, cell wall modification, and active pathogen sensing and signaling receptors. Such features could facilitate early protective responses that limit nematode invasion or establishment and influence the timing and magnitude of subsequent inducible defenses (Hückelhoven and Schouten, 2024; Kumar et al., 2021; Tomar et al., 2026). In soybean cultivars, distinct constitutive gene and/or metabolite expression patterns have been observed between resistant genotypes (Chu et al., 2022; Klink et al., 2011) and/or susceptible genotypes, with resistant responses generally showing greater constitutive expression (Cook et al., 2012). Moreover, in other plant systems, resistant genotypes showed greater constitutive abundance of specialized metabolites, such as phenolics and alkaloids, compared to susceptible genotypes prior to nematode infection (Hauri et al., 2024; Rahaman et al., 2020). Here, S54 exhibited higher constitutive abundance of selected defense-, redox-, and metabolism-associated features under control conditions (**Fig. 3**) while both genotypes subsequently followed different molecular trajectories after infection (**Fig. 2**). In wild soybean, SCN resistance may be conferred by a combination of constitutive molecular preparedness and stronger induced defense responses that enable a rapid and effective response to infection. Interestingly, wild soybean-based resistance cannot solely be attributed to strong expression of major SCN resistance mechanisms characterized in cultivated soybean (Miraeiz *et al*., 2020). From our previous study using Race 5 SCN (HG type 2.5.7), the resistant wild soybean genotype NRS100 did not show significant infection-induced expression of genes associated with Peking-type or PI88788-type resistance (Kofsky *et al*., 2021). Similarly, in the present study, we found limited evidence of strong constitutive expression or induction of selected genes associated with established cultivated-soybean resistance mechanisms, including SNAP proteins (Haarith et al., 2025), wound-inducible domain proteins (Dong and Hudson, 2022), serine hydroxymethyltransferases (Zhang et al., 2026), and amino acid transporters (Du et al., 2025) in either S54 or S67. Together, these observations suggest that SCN resistance in S54 may employ additional defense-associated processes beyond those commonly emphasized in cultivated soybean.

### Resistance and susceptibility are associated with distinct gene co-expression networks

To examine gene co-expression patterns associated with S54 and S67 responses to SCN infection, we conducted weighted gene co-expression network analysis (WGCNA) using control and SCN-treated samples across timepoints. Our WGCNA analysis identified 18 co-expression modules and subsequently assigned functional categories based on associated GO/KEGG enrichment terms. Module-trait correlations were evaluated to identify modules associated with genotype, SCN treatment, or timepoint (**Fig. 4A**).

**Figure 4.**
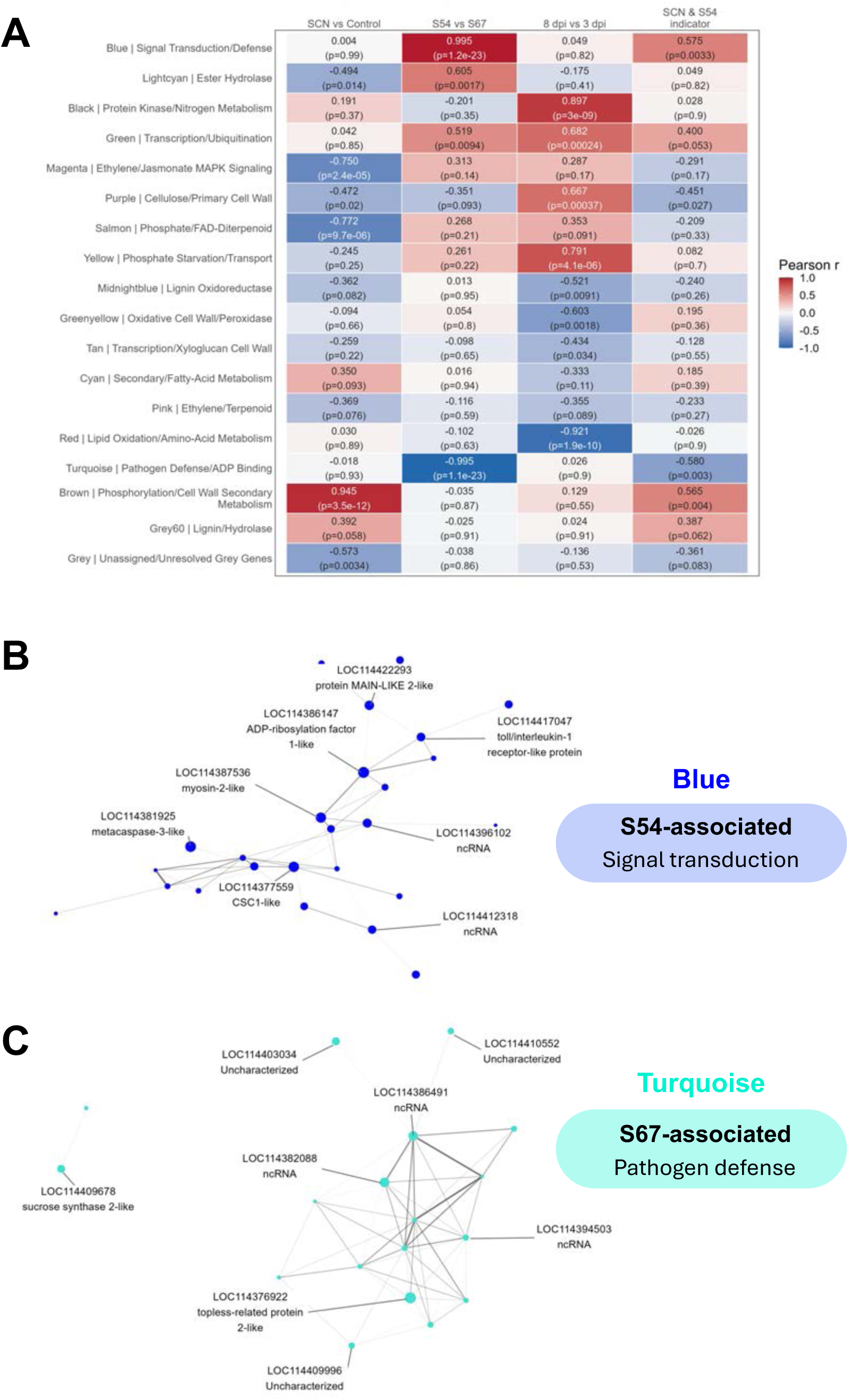
**Gene co-expression analysis**. WGCNA modules with gene enrichment-based assigned functional categories. (A) Module-trait heatmap (n = 24, control and SCN-treated) shows the relationship strength based on Pearson’s correlation (r). (B) S54-associated, signal transduction-related module network (Blue) of top co-expressed genes. (C) S67-associated, pathogen defense-related module network (Turquoise) of top co-expressed genes.

Among the identified modules, genotype-associated (S54 vs. S67) modules comprised the largest gene counts across control and SCN-treated samples and exhibited the strongest correlations regardless of SCN infection. Genes within the Blue module (“signal transduction,” n = 1,223) were highly expressed across control and SCN-treated samples in S54 and showed the strongest positive association with S54 (r = 0.99) relative to S67 (**Fig. 4B**). In contrast, genes within the Turquoise module (“pathogen defense,” n = 1,315) were highly expressed across control and SCN-treated samples in S67 and negatively correlated with S54 (r = -0.99) relative to S67 (**Fig. 4C**). For these genotype-associated modules, we generated individual WGCNA networks for the Blue (“signal transduction”) and Turquoise (“pathogen defense”) modules and identified top-ranked hub genes based on network connectivity and kME values (**Fig. 4B–C**). These results suggest that resistant and susceptible genotypes are associated with distinct gene co-expression patterns that are consistent across control and SCN-treated samples, thereby providing evidence for distinct constitutive expression patterns between genotypes.

The association between S54 and the Blue-signal transduction module is consistent with our other evidence suggesting that signaling-related responses are prominently associated with resistance. Although this module was primarily associated with genotype rather than SCN treatment, its functional enrichment, together with the differential expression of signaling-related genes following infection, supports a potential role for these processes in S54 resistance. GO enrichment analysis identified several upregulated signal transduction-related processes in S54 during SCN infection, including hormone-mediated signaling, signaling receptor activity, and glutamate-gated receptor activity (**Fig. S1**). Consistent with other work in resistant wild soybean (Miraeiz *et al*., 2020), S54 also showed a greater abundance and stronger induction of several receptor-like kinases (*RLKs*) and receptor-like proteins (*RLPs*), as well as wall-associated kinases (*WAKs*), chitin elicitor receptor kinase 1 (*CERK1*), and leaf rust 10 disease resistance locus RLKs. These pattern recognition receptors are critical for early sensing and detection of pathogen- and damage-associated molecular patterns (PAMPs and DAMPs, respectively) and initiating targeted plant immune responses coordinated between PAMP-triggered immunity (PTI) and effector-triggered immunity (ETI) signaling pathways (Kumar *et al*., 2021).

Following pathogen perception, signaling molecules such as calcium, ROS, and ncRNAs can transduce signals that activate downstream defense pathways (Ding et al., 2022). In S54, we observed stronger induction of signal molecule-mediated activity, especially calcium-activated signaling components such as calcium receptor proteins (e.g., *CaMs*, *CDPKs*) and calcium transporters (e.g., *GLRs*, *CNGCs*) during early SCN infection. Rapid changes in localized calcium concentrations can trigger a cascade of cellular events and defense signaling pathways (Ding *et al*., 2022). Notably, glutamate receptor channels (*GLRs*) can modulate calcium fluxes and activate downstream components of calcium-mediated defense pathways (Saberi Riseh et al., 2026). S54 also showed strong induction (>1,000-fold change) of a calcium-activated phospholipase A1-ligamma gene, which modulates calcium signaling and is involved in lipid-mediated signaling, lipid metabolism, membrane trafficking, and oxylipin biosynthesis (Ryu, 2004; Xing et al., 2021). Collectively, our WGCNA, differential expression, and enrichment findings support the association between signal transduction processes and the resistant response of S54. Overall, these co-expression patterns may provide a framework for identifying hub genes that underpin the contrasting genotypic responses to SCN infection.

Interestingly, the Turquoise module, enriched for pathogen defense-related functions, was strongly associated with susceptible S67, whereas the signal transduction-related Blue module was associated with resistant S54 (Fig. 4). This contrast highlights that the abundance of defense-associated transcripts does not necessarily translate into effective resistance. Although both genotypes exhibited SCN-responsive molecular changes, S54 showed more extensive transcriptional responses at 3 dpi, whereas S67 exhibited more DEGs at 8 dpi (Fig. 2). The greater transcriptional response in S67 at 8 dpi may reflect continued infection-associated stress or compensatory defense responses that are insufficient to restrict SCN infection. In contrast, the constitutive abundance of selected defense-associated features and distinct early responses in S54 may contribute to more effective resistance. Together, these observations suggest that resistance may depend not simply on the magnitude of defense-associated responses, but also on their molecular composition and temporal regulation.

### Multi-omics integration reveals distinct genotype- and timepoint-associated networks

To integrate our transcriptomic, metabolomic, and proteomic datasets and identify correlated molecular features associated with genotype and temporal responses to SCN infection, we applied DIABLO analysis to construct multi-omics networks and correlative heatmaps (**Fig. 5, Fig. S2**). Our DIABLO networks identified the strongest cross-omics pattern of correlated genes, metabolites, and proteins that best explains the biological variance between sample groups. From this analysis, loading values represent the weighted contribution of each feature, where the higher loading value is more influential. These networks included selected features (top 5 per omics block) represented across genotype-associated networks (**Fig. 5**) and timepoint-associated networks (**Fig. S2**).

**Figure 5.**
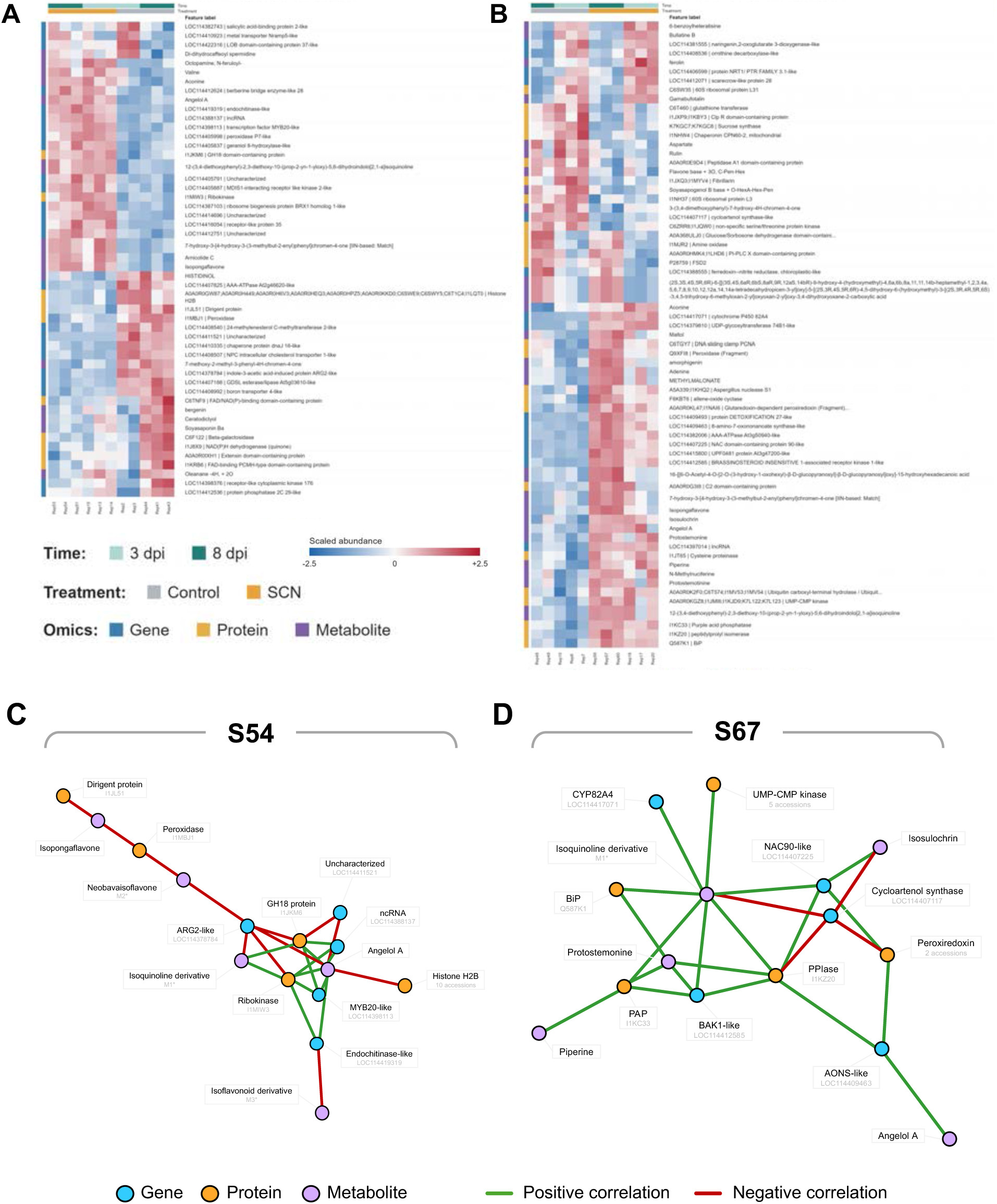
**DIABLO network analysis**. Heatmaps show expression (scaled abundance) of all DIABLO-selected features across timepoints and treatments for (A) S54 and (B) S67. Genotype-associated DIABLO cross-omics networks (SCN versus control, 3 and 8 dpi pooled) show the top 5 correlated features per omics block associated with (C) S54 and (D) S67 responses. Correlated features are represented by colored nodes and connecting lines. Full compound names for select network terms: M1 = 12-(3,4-diethoxyphenyl)-2,3-diethoxy-10-(prop-2-yn-1-yloxy)-5,6-dihydroindolo[2,1-a]isoquinoline. M2 = 7-hydroxy-3-[4-hydroxy-3-(3-methylbut-2-enyl)phenyl]chromen-4-one. M3 = 7-methoxy-2-methyl-3-phenyl-4H-chromen-4-one.

In our S54-associated analysis, the top DIABLO loadings included defense-, regulatory-, redox-, and specialized metabolism-associated genes and proteins including chitinases (endochitinase and GH18), a *MYB20* transcription factor, ncRNA, dirigent protein, and peroxidase. Notably, specialized defense metabolites associated with SCN infection in this network included phenolics, alkaloids, and most prominently, isoflavonoids (**Fig. 5A, 5C**). In our S67-associated analysis, the top DIABLO loadings included defense-, signaling-, detoxification-, and metabolism-associated features including a cytochrome P450 CYP82A4, a brassinosteroid-insensitive 1-associated receptor kinase 1 (*BAK1*), an 8-amino-7-oxononanoate synthase (*AONS*), a NAC90 transcription factor, purple acid phosphatase, a glutaredoxin-dependent peroxiredoxin, and specialized metabolites that were largely distinct from those in S54 (**Fig. 5B, 5D**). Notably, both genotype-associated networks highlight the prevalence of specialized metabolites associated with plant chemical defense, however the composition of these metabolites is distinct between genotypes. In particular, S54’s top 5 network contained multiple isoflavonoid compounds, that were not selected in S67’s top 5 network (**Fig. 5C–D**).

These differences highlight the role of defense-associated metabolites in supporting plant resistance to pathogens such as SCN. Flavonoids and isoflavonoids derived from phenylpropanoid metabolism have been implicated in plant-nematode interactions, such as soybean-SCN, resulting in altered nematode chemotaxis, chemoreception, penetration, development, and survival (Chin et al., 2018; Yang and Wang, 2024; Yang et al., 2024). Recently, Jiang et al. (2025) identified a UDP-glucosyltransferase encoding gene through multi-omics analysis and demonstrated that overexpression of this gene increased isoflavonoid accumulation, thereby enhancing SCN resistance (Jiang et al., 2025). MYB transcription factors are versatile regulators of specialized metabolism pathways including phenylpropanoids such as isoflavonoids (Mariyam et al., 2025; Pratyusha and Sarada, 2022; Yang and Wang, 2024). Given the S54-associated network selection of *MYB20* together with multiple isoflavonoids, *MYB20* may be involved in regulating isoflavonoid accumulation in S54. In conjunction with the direct presence of defense metabolites in S54, we observed indirect evidence of defense-associated metabolism. Notably, higher induction of beta-glucosidase genes in S54 may have contributed to the release of defense metabolites and signaling molecules that were previously “inactive” (e.g., glycosides). In a recent study, overexpression of a beta-glucosidase gene enhanced SCN resistance in a soybean cultivar (HN531) by hindering SCN development post-penetration (You et al., 2025). Collectively, our DIABLO analysis revealed genotype-specific and timepoint-specific networks of correlated genes, proteins, and metabolites, with defense- and signaling-associated features most prominently associated with S54’s response.

### Integrated DIABLO and WGCNA analyses prioritize candidate defense genes

To identify potential candidate genes supported by both DIABLO and WGCNA analyses, DIABLO-selected genes were mapped to WGCNA modules to examine shared molecular patterns. Among these overlapping genes, candidates were prioritized based on DIABLO ranking, kME values, and WGCNA rankings. Expression profiles of DIABLO-selected genes and top WGCNA hubs and their associated modules were visualized in a heatmap (**Fig. 6A**). These candidate genes revealed distinct constitutive and induced response differences between genotypes across timepoints (**Fig. 6A**). Across the WGCNA modules, several genes were also identified in the DIABLO genotype-associated networks (Fig. 5C–D) and timepoint-associated networks (Fig. S2). Genes of interest included a putative steroid-binding protein 3 (*SBP3*), pleiotropic drug resistance protein 3 (*PDR3*), cysteine-rich receptor-like protein kinase 6 (*CRK6*), endochitinase, an uncharacterized gene (LOC114411280), and a ncRNA. Other notable genes not identified in our DIABLO networks but present in our WGCNA modules included a TMV resistance protein N-like gene, a receptor-like protein 33, WRKY70, and other defense- and signaling-related genes (**Fig. S3**).

**Figure 6.**
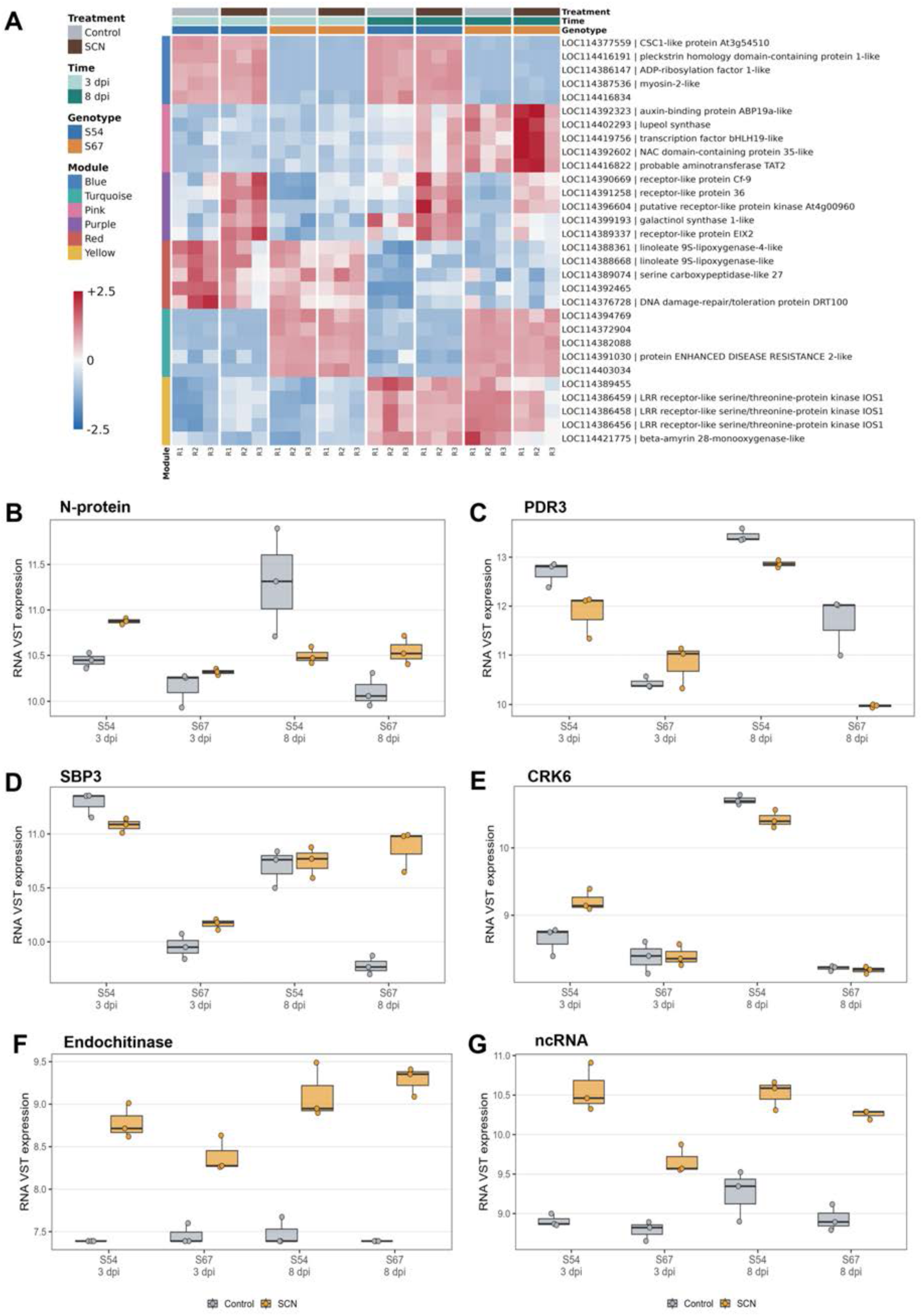
**DIABLO-WGCNA selected hubs and priority gene candidate expression**. (A) Heatmaps show WGCNA hub gene (top 5) expression (z-score) and associated WGCNA modules (treatments and timepoints pooled). Boxplots show the expression (VST) of key DIABLO network-selected gene candidates. (B) TMV resistance protein N-like (N-protein). (C) pleiotropic drug resistance 3-like (PDR3). (D) probable steroid binding protein 3 (SBP3). (E) cysteine-rich receptor-like protein kinase 6 (CRK6). (F) endochitinase-like. (G) ncRNA.

The TMV resistance protein N-like gene and PDR3 gene showed higher constitutive expression in S54 than in S67, especially at 8 dpi (**Fig. 6B–C**). SBP3 and CRK6 maintained higher expression in S54 across control and SCN-treated conditions relative to S67 (**Fig. 6D–E**). In contrast, the endochitinase and ncRNA genes showed greater induction in S54 compared to S67, mainly at 3 dpi (**Fig. 6F– G**). Together, these expression profiles highlight distinct patterns of constitutive and SCN-induced responses among key genes from our DIABLO and/or WGCNA analyses. Collectively, several of these genes have functions consistent with roles in plant defense and signaling. TMV resistance N-like proteins belong to the broader NLR immune-receptor class, whose members function as intracellular pathogen sensors and/or downstream immune-signaling components (Locci and Parker, 2024). Similarly, cysteine-rich receptor-like kinases (*CRKs*) participate in cell-surface signaling and have emerging roles in pattern-triggered immunity, ROS-associated signaling, and other plant stress responses (Krasensky-Wrzaczek and Wrzaczek, 2024). PDR proteins are members of the plant ABCG transporter family, which can mediate transport and/or secretion of specialized metabolites, thereby contributing to chemical defense.

In plants, chitinases, classified as pathogenesis-related proteins, can participate in defense responses regulated by hormones and transcriptional regulators, including *WRKY* transcription factors and ncRNAs. Endochitinases hydrolyze internal glycosidic bonds within chitin polymers and have established roles in plant defense against chitin-containing pathogens. Although the direct contribution of chitinase activity to SCN resistance remains unclear, our integrated analyses identified an endochitinase gene and a GH18 chitinase-related protein among the S54-associated molecular features. Notably, the endochitinase transcript exhibited stronger induction in resistant S54 at 3 dpi than in susceptible S67. Together, these findings highlight chitinase-associated responses as promising candidates for further investigation of natural SCN resistance in wild soybean (Mahajan et al., 2025; Vaghela et al., 2022). Various ncRNAs were observed within our WGCNA and DIABLO genotype-associated networks. While ncRNAs (e.g., miRNAs) do not encode any proteins, they can actively regulate plant innate immunity and stress responses by modifying or inhibiting the translation of gene transcripts such as R genes, transcription factors, and signaling receptors (Boccara et al., 2014; Deng et al., 2018; Ding *et al*., 2022). Notably, in cultivated soybean, miRNAs have been reported to increase isoflavonoid content (Gupta et al., 2019) and enhance SCN resistance by targeting the expression of *MYB* transcription factors (Lei et al., 2021). Although the specific roles of ncRNAs identified in our study remain unclear, previous studies suggest that ncRNAs can modulate transcription factor expression (e.g., *MYBs*) and regulate broader defense networks (Mariyam *et al*., 2025).

Integration of our WGCNA and DIABLO analyses narrowed our broader multi-omics findings to a set of high-priority candidate genes exhibiting distinct constitutive and SCN-induced expression patterns. These patterns include higher basal expression in S54, sustained expression in S54, and stronger induced responses in S54 at 3 and/or 8 dpi. Overall, these patterns are consistent with the broader differences observed between resistant (S54) and susceptible (S67) responses across our multi-omics datasets and suggest that both constitutive and rapidly induced defense responses may contribute to SCN resistance in S54. These candidate genes provide promising targets for future functional studies to determine their contributions to SCN resistance.

### Summary and Perspectives

Here, we applied a systems-level, integrated multi-omics approach to characterize transcriptomic, metabolomic, and proteomic responses to SCN infection in resistant (S54) and susceptible (S67) wild soybean genotypes. Across early and late timepoints, both genotypes shared many core SCN-responsive processes but differed in their constitutive and induced responses, gene co-expression networks, and correlated multi-omics features. Notably, S54 exhibited higher constitutive abundance of selected defense-associated features and distinct early infection responses, whereas S67 showed more extensive transcriptional changes at 8 dpi. These findings suggest that effective SCN resistance may depend not simply on the magnitude of defense responses, but also on constitutive molecular preparedness and the composition and timing of induced responses. Collectively, these analyses identify molecular features associated with SCN resistance and susceptibility and highlight potential candidate genes and biological pathways for future functional investigation. Our findings suggest that S54 may be able to more effectively sense pathogenic invasion and subsequently respond to infection more quickly than S67, thereby contributing to its resistance.

Beyond advancing our understanding of SCN resistance in wild soybean, this study highlights the broader value of crop wild relatives as systems for investigating naturally evolved resistance mechanisms and identifying promising genetic resources for crop improvement. Our findings also raise new questions and research priorities for future investigations. Although we sampled two distinct timepoints, these represent only a limited window of the dynamic host response to SCN infection, from initial penetration to feeding-site establishment and nematode development. Expanded temporal sampling will further elucidate the timing and progression of genotype-specific responses and their relationship to constitutive molecular differences. Furthermore, candidate genes prioritized through individual omics, WGCNA, and integrated multi-omics analyses provide opportunities for functional validation and hypothesis testing. Finally, extending these analyses across additional wild soybean genotypes and SCN populations will help determine which features are specific to the S54–S67 comparison and which represent broader SCN resistance mechanisms.

Ultimately, a more holistic view of SCN resistance will deepen our understanding of the molecular dynamics shaping soybean-SCN interactions. Resolving complex interactions within plant-pathogen systems requires large-scale gene discovery efforts and may be accelerated by emerging multi-omics, single-cell, spatial omics, and computational tools, alongside conventional breeding and functional genomics. In particular, single-cell approaches could help resolve cell-type-specific responses associated with nematode penetration and feeding-site establishment in infected root tissues. Together, these approaches could provide a more comprehensive understanding of SCN resistance mechanisms, beyond those characterized in cultivated soybean, while identifying promising molecular targets for functional validation and the development of improved SCN-resistant cultivars.

## AUTHOR CONTRIBUTIONS

B-H.S. conceived and designed the project. P.Y. conducted the experiment. All authors contributed to project discussions. B-H.S. and P.Y. collected transcriptomics data. S.C., S.M., and Y.S. collected metabolomics and proteomics data. M.S.M., Y.W., and T.Z. performed integrative analyses. T.Z. and P.Y. generated manuscript figures. S.M. and T.Z. wrote the data analysis sections. P.Y. wrote the manuscript draft. P.Y., B-H.S., S.C., S.M., and T.Z. revised the manuscript. All authors approved the final manuscript.

