## Supplemental Figures for "Time-resolved multi-omics reveals constitutive defense preparedness and early responses to soybean cyst nematode in wild soybean"

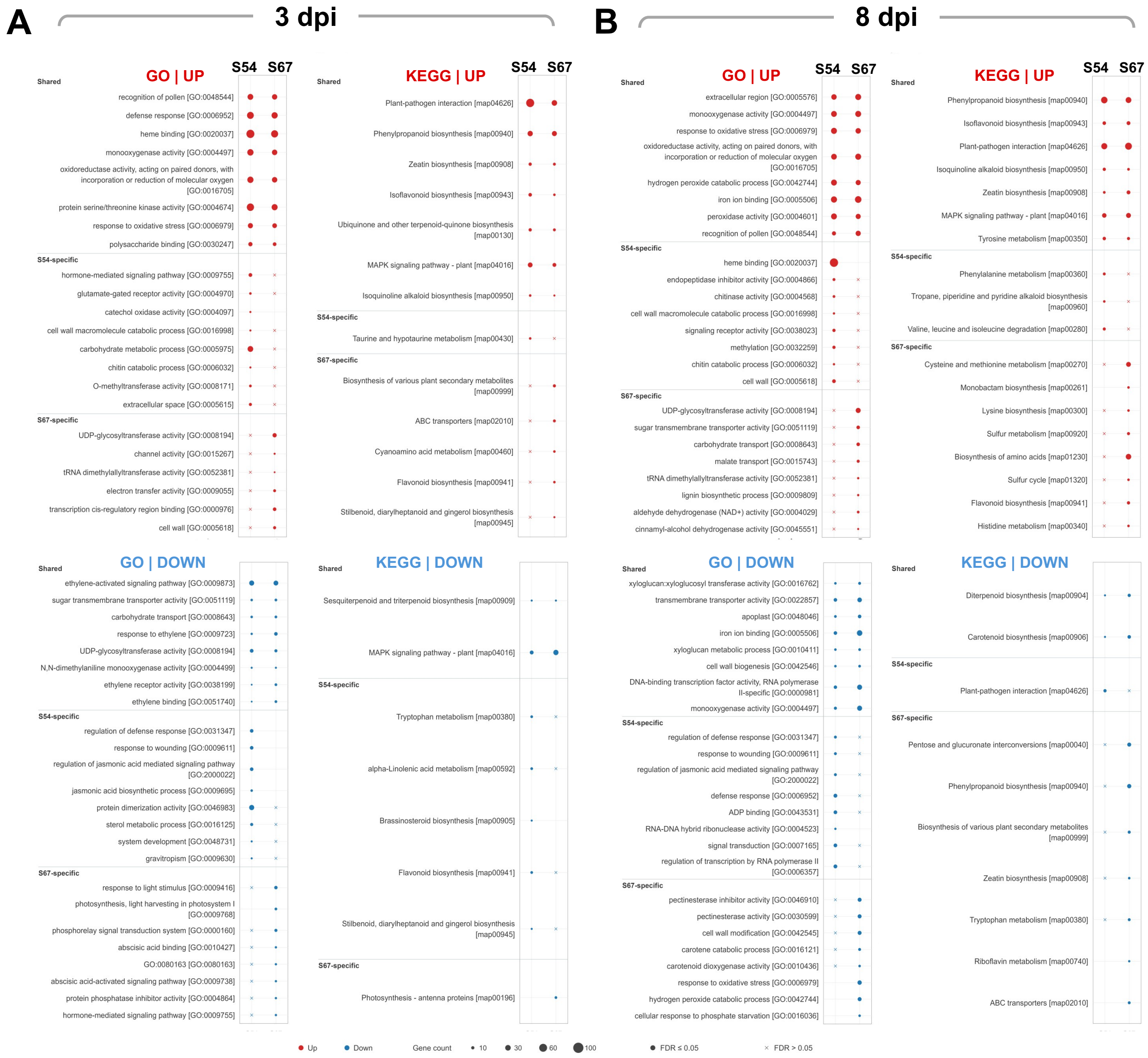


**Figure S1**. **GO and KEGG enrichment analysis.** Dot plots showing shared and genotype-specific GO and KEGG enrichment terms for upregulated (Red) and downregulated (Blue) DEGs at (A) 3 dpi and (B) 8 dpi. Dots = FDR ≤ 0.05.

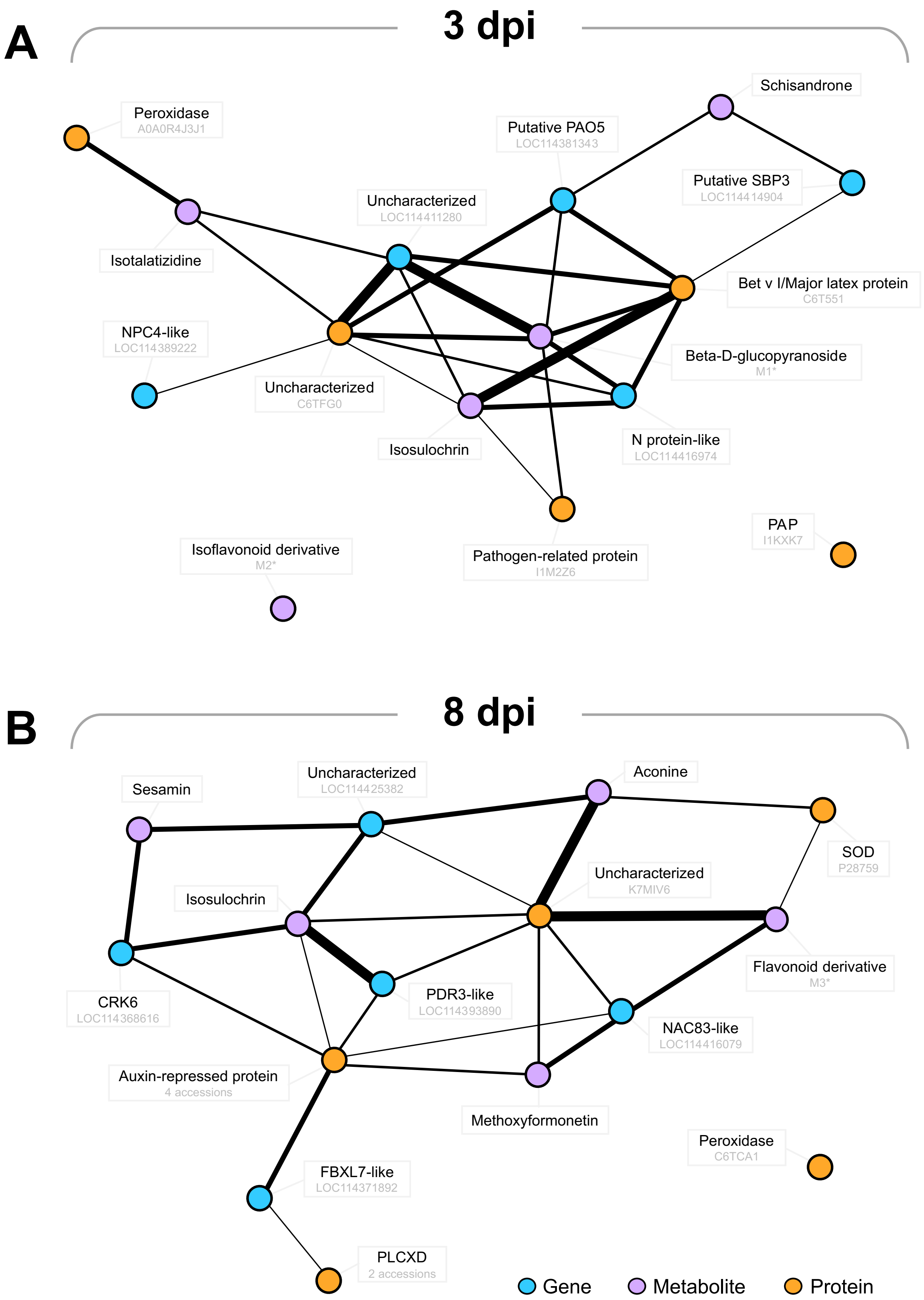
A
3 dpi
Peroxidase
A0A0R4J3J1
Schisandrone
Putative PAO5
LOC114381343
Putative SBP3
LOC114414904
Uncharacterized
LOC114411280
Isotalatizidine
Bet v I/Major latex protein
C6T551
NPC4-like
LOC114389222
Beta-D-glucopyranoside
M1*
Uncharacterized
C6TFG0
Isosulochrin
N protein-like
LOC114416974
PAP
I1KXK7
Isoflavonoid derivative
M2*
Pathogen-related protein
I1M2Z6
B
8 dpi
Sesamin
Uncharacterized
LOC114425382
Aconine
SOD
P28759
Uncharacterized
K7MIV6
Isosulochrin
Flavonoid derivative
M3*
CRK6
LOC114368616
PDR3-like
LOC114393890
NAC83-like
LOC114416079
Auxin-repressed protein
4 accessions
Methoxyformonetin
Peroxidase
C6TCA1
FBXL7-like
LOC114371892
PLCXD
2 accessions
Gene
Metabolite
Protein

**Figure S2**. **Timepoint-associated DIABLO cross-omics networks**. Networks (SCN only, genotypes pooled) show the top 5 correlated features per omics block associated with (A) 3 dpi and (B) 8 dpi responses. Correlated features are represented by colored nodes and connecting lines. Thicker lines signify a stronger correlation. Full compound names for select network terms: M1 = 2-hydroxycyclohexyl 2-O-[(2E)-3-(3,4-dihydroxyphenyl)-2-propenoyl]-β-D-glucopyranoside. M2 = isoflavanone base + 2O, 1MeO, 2Prenyl. M3 = flavonol base + 2MeO.

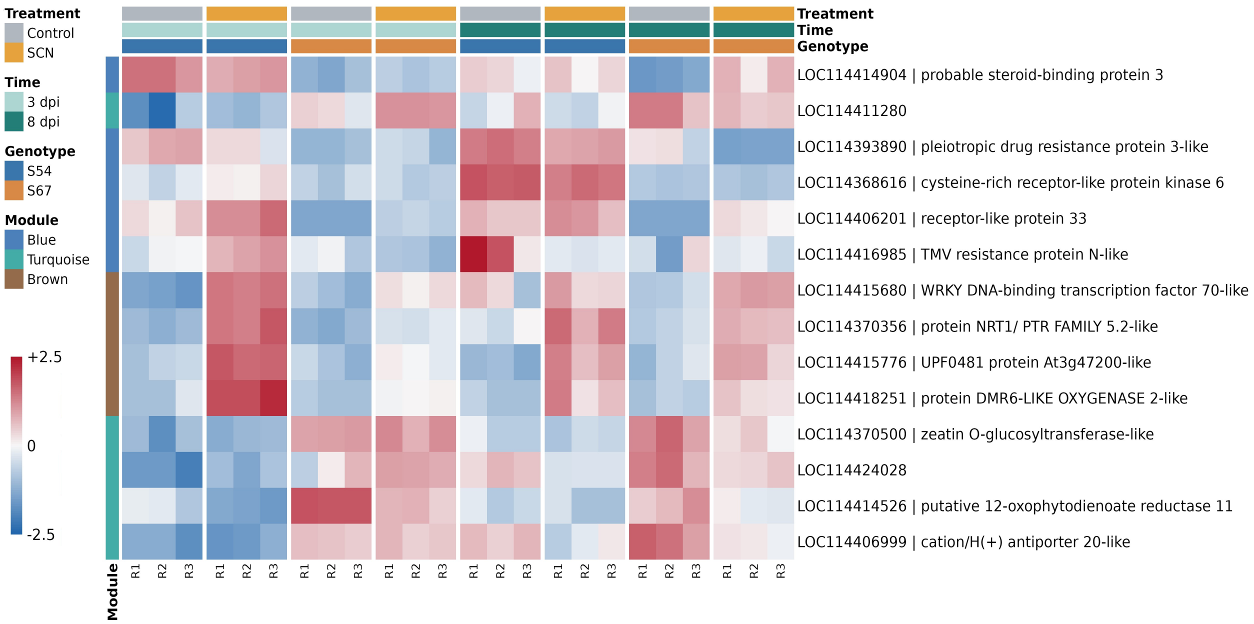


**Figure S3**. **DIABLO-WGNCA selected gene candidate expression**. Heatmap showing expression (z-score) of DIABLO-selected, WGCNA-assigned gene candidates. 3 dpi-associated DIABLO network genes: Uncharacterized (LOC114411280), probable steroid binding protein 3 (SBP3) (LOC114414904). 8 dpi-associated DIABLO network genes: pleiotropic drug resistance protein 3-like (PDR3) (LOC114393890), cysteine-rich receptor-like protein kinase 6 (CRK6) (LOC114368616).
